# Plasmonic Stimulation of Gold Nanorods for the Photothermal Control of Engineered Living Materials

**DOI:** 10.1101/2022.11.30.518571

**Authors:** Selim Basaran, Sourik Dey, Shardul Bhusari, Shrikrishnan Sankaran, Tobias Kraus

## Abstract

Engineered living materials (ELMs) use encapsulated microorganisms within polymeric matrices for biosensing, drug delivery, capturing viruses, and bioremediation. It is often desirable to control their function remotely and in real time. Suitable, genetically engineered microorganisms respond to changes of their environment. Here, we combine this local sensitivity with a nanostructured encapsulation material to sensitize the ELM for infrared light. Previously, blue light has been used to stimulate microorganisms that contain optogenetic modules responsive to those wavelengths without the need for exogenous cofactors. Here, we use plasmonic gold nanorods (AuNR) that have a strong absorption maximum at 808 nm, a wavelength where human tissue is relatively transparent. Biocompatible composites of a Pluronic-based hydrogel and AuNR are prepared without agglomeration; they react to illumination by local heating. We measure a photothermal conversion efficiency of 47 % in transient temperature measurements. Steady-state temperature profiles from local photothermal heating are quantified using infrared photothermal imaging, correlated with measurements inside the gel, and applied to stimulate thermoresponsive bacteria. Using a bilayer ELM construct with the thermoresponsive bacteria and the thermoplasmonic composite gel in two separate but connected hydrogel layers, it is shown that the bacteria can be stimulated to produce a fluorescent protein using infrared light in a spatially controlled manner.

## 1. Introduction

Engineered living materials (ELMs) are a rapidly developing class of composite materials born from the combination of synthetic biology and materials science.[1, 2] In ELMs, living microorganisms (bacteria, yeast, algae etc.) are combined with inanimate materials to yield advanced constructs applicable as biosensors,[3] self-healing adhesives,[4] biofilters to trap metals and viruses,[5, 6] smart drug delivery devices,[7] soft robots,[8] and other devices. In most of these cases, the functionalities are realized through genetic engineering of the microorganisms while the material component acts as a supportive scaffold.

Light is an attractive stimulus for ELMs that can be applied at different intensities, wavelengths, and spatial dimensions at low cost. Light has been used for patterning,[9, 10] actuation,[11] and drug release[7] in ELMs. Existing systems typically use blue light with short wavelengths that act on optogenetic systems in bacteria.[7, 10] Blue light, however, exhibits low penetration depths *in vivo*, limiting e.g. translational potential for implanted ELMs.[12] Green-, red- and near-infrared (NIR)-responsive optogenetic circuits[9] have been reported but require co-factors that are not naturally present in most bacteria and therefore need to be externally supplied or produced by the bacteria themselves through engineered complex enzyme cascades.

Here, we follow an alternative strategy that combines an inanimate functional material which acts as an optical absorber with heat-responsive genetic circuits. The combination retains the advantages of long-wavelength light (spatiotemporal control with relatively high penetration depth in skin) without requiring light-responsive genetic circuits. The optical absorber is based on gold nanorods (AuNR) that are well-accepted in biomedical applications due to their biocompatibility,[13, 14] unique absorption properties in the visible and NIR regions arising from conduction electron-light interactions,[15, 16] localized surface plasmon resonance (LSPR) that can be tuned by changing the geometry,[17] and high photothermal efficiency.[18] Nanorods have thus been used in cancer therapy,[19-21] drug delivery,[20, 22-24] and photothermal therapy (PTT).[18] They can trigger hyperthermia in tissues [25, 26] and induce cell death via necrosis and apoptosis at temperatures above 45°C.[27] Gold nanorods of suitable geometry can efficiently absorb NIR light in the range of the tissue transparency window of 650–1350 nm.[28, 29] Stimulation at such wavelengths can be safe to tissues, cells, and bacteria with the ability of deep tissue penetration.[12, 30, 31] The absorption of NIR light causes a plasmon oscillation of the conduction electrons that is dampened by lattice collision resulting in thermalization, which heats the surrounding matrix.[32]

It is challenging to use free, dispersed AuNR in the body. Challenges include insufficient retention [30] and the need for specific surface functionalization for targeting[26, 28] and to prevent cytotoxicity.[33-35] Here, we avoid such problems by embedding the Au NR in hydrogels, resulting in gold nanorod nanocomposite hydrogels (GNC) that retain the Au NR and enable spatiotemporal control of stimulation by local illumination.[36] Such composites have been applied for PTT by local heating,[34] for drug delivery by shrinking or decomposing the gel through the generated heat,[30, 36-39] and photodynamic treatments with dyes that generate radical oxygen species.[26, 28] Comparable GNC hydrogels have been formulated to mimic the mechanical properties of human tissues and minimize immune reactions.[40] They can be injected locally and retain the Au NR at the treatment site.[41]

Local heating by several 10 °C is possible with GNCs.[42] A temperature increase to 45°C or above has been demonstrated and used to kill cancer cells.[43] Here, we use moderate temperature increases to activate bacteria in ELMs. They enable a more targeted approach to a range of therapies, where moderate local heating induces the release of active components from bacteria.[1] The encapsulation in ELM enables photothermal interaction of Au NR and bacteria while physically separating the genetically modified bacteria from the organism.[44]

In this study, we demonstrate the activation of thermogenetically engineered bacteria in an ELM that we sensitize by a GNC to make it NIR-responsive. A bilayer hydrogel structure combines a bottom layer as ELM with a GNC top layer. We quantify and control the thermoplasmonic conversion to reach temperature levels that are harmless for the surrounding tissue and optimal for the activation of the bacteria. Activation is then quantified via fluorescence microscopy using bacteria that express the fluorescent protein mCherry.

## 2. Materials and Methods

### 2.1 Materials

Gold nanorods (length: 41 nm, diameter: 10 nm) capped with CTAB in water were purchased from Nanopartz Inc. (Canada). The hydrogel precursor, Pluronic diacrylate (PLUDA), was synthesized by reacting Pluronic F127 (Plu, MW≈12600 g/mol, Sigma-Aldrich) with acryloyl chloride in the presence of triethylamine according to a reported protocol.[45] 30% (w/v) solutions were prepared in milliQ water (PLUDA_MQ_) and LB medium (PLUDA_LB_) containing Irgacure 2959 and photoinitiator at 0.2 % w/v.

Microscopy slides with 15 wells (type “μ-slide Angiogenesis”) with uncoated bottoms were obtained from ibidi (Gräfelfing, Germany). Silicone oil (350 cSt, Sigma Aldrich) was used to cover the ELM and prevent it from drying.

### 2.2 Creation of thermoresponsive mCherry ClearColi Strain

The mCherry double-stranded gene insert was procured from Integrated DNA Technologies (Coralville, USA) and assembled into the pTlpA39-Wasabi vector (Addgene #86116)[46] using the NEBuilder HiFi DNA Assembly Cloning Kit (New England Biolabs, GmbH). The mCherry gene was inserted downstream of the PtlpA promoter by replacing the mWasabi reporter gene. The recombinant plasmid construct was sequence verified (Eurofins GmbH, Germany) and subsequently transformed in the ClearColi BL21(DE3) strain (BioCat GmbH, Germany) and maintained in LB Miller medium (Carl Roth GmbH, Germany) supplemented with 100 μg/mL Ampicillin.

### 2.3 Preparation and characterization of GNC hydrogel

The CTAB stabilized gold nanorods were centrifuged at 12,500 rpm for 8 min at room temperature and the surfactant was removed by pipetting. Then the suspension was diluted with the PLUDA_MQ_ to the desired concentrations at 4°C. The mixture was incubated at room temperature for 5 min to allow gelation. The GNC hydrogel was exposed to a UV lamp (365 nm, 6 mW/cm^2^) inside Alpha Innotech FluorChem Q system (Biozym, Oldendorf, Germany) for 2 min to crosslink the diacrylate groups. The GNC hydrogel was characterized by UV-VIS-NIR spectrometer. The absorbance was measured in a 10 mm disposal cuvette. The baseline was pure hydrogel in cuvette.

The distribution of the gold nanorods was observed by cryo-TEM. A small amount of the GNC hydrogel was placed on a holey carbon grid (Plano, Wetzlar, Germany, type S1474-4), blotted for 2 s, and plunged into liquid ethane using a Gatan (Pleasanton, CA, United States) CP3 cryo plunger operating at *T* = -165 °C. The vitrified sample was transferred to a Gatan model 914 cryo-TEM sample holder and investigated by bright field TEM (JEOL, Akishima, Japan, JEM-2100 LaB6) imaging at 200 kV accelerating voltage and *T* = -170 °C under low-dose conditions. A Gatan Orius SC1000 CCD camera was used for image acquisition (2 s exposure time).

### 2.4 Preparation of bilayer hydrogel structures

The hydrogel precursor for non-bacterial experiments contained 30 % (w/v) of a diacrylated Pluronic F127 (PLUDA) in Milli-Q water, and 0.2 % of the photoinitiator Irgacure (2-benzyl-2-(dimethylamino)-1-[4-(morpholinyl) phenyl)]-1-butanone). A dispersion of gold nanorods (AuNR@CTAB) with an Optical Density (OD) of OD_808nm_ = 40 was obtained by centrifugation of a dilute suspension at 12,500 rpm for 8 min at room temperature. It was immediately mixed with PLUDA_MQ_ (180 μL) at 4 °C to dilute to the desired OD_808nm_ = 4. The mixture was immediately used to prepare GNCs. The hydrogel precursor for the bacterial hydrogels contained 30 % (w/v) of PLUDA in lysogeny broth (LB) medium, and 0.2 % of the photoinitiator Irgacure. This mixture is denoted as PLUDA_LB_ in the following.

The thermoresponsive mCherry producing ClearColi BL21(DE3) strain was inoculated in LB Miller – 2% NaCl medium supplemented with 100 μg/mL ampicillin and incubated at 37 °C, 250 rpm shaking conditions for 18 h. The bacterial culture reached the late logarithmic phase and the bacterial cell density was adjusted to OD_600nm_ = 2 by centrifugation (9,600 rpm, 4 °C, 3 min). Determination of the cell density was done by a NanoDrop Microvolume UV-Vis 175 spectrophotometer (ThermoFisher Scientific GmbH, Germany) and resuspension in fresh medium was done by vortexing for less than 1 min. A volume of 10 μL was dispersed in PLUDA_LB_ (90 μL) at 4 °C and homogenized to a resulting OD_600nm_ = 0.2. The mixture was immediately used to prepare the bacterial hydrogel layers.

The bilayer hydrogel structures were prepared in patterned polymer microscopy slides with 5 × 3 microwells and uncoated bottoms (Fig. 1). Each microwell had an inner well (10 μL, diameter 4 mm) that was used to form the bacterial hydrogel layer and an upper well (50 μL, diameter 5 mm) that was used to hold the GNC hydrogel layer. 10 μL of the bacterial hydrogel precursor solution were pipetted at 4°C in the lower well and then incubated for 5 min at room temperature for gelation. The physically assembled gel was photopolymerized at a wavelength of 395 nm and a power density of 6 mW/cm^2^ for two minutes using a FluorChemQ light source (Alpha Innotech). The GNC hydrogel precursor (20 μL) was injected in the upper well, incubated for 5 min at room temperature for physical gelation, and UV cured in the same way as the lower gel. The obtained bilayer hydrogel structure was topped with 20 μL silicone oil to prevent drying. The result was a cylindrical shaped bacterial hydrogel with a diameter of 4 mm and a height 1 mm and a cylindrical GNC hydrogel with a diameter of 5 mm and a height of 1 mm. A 1 mm thick silicone layer covered it to prevent drying.

**Figure 1:**
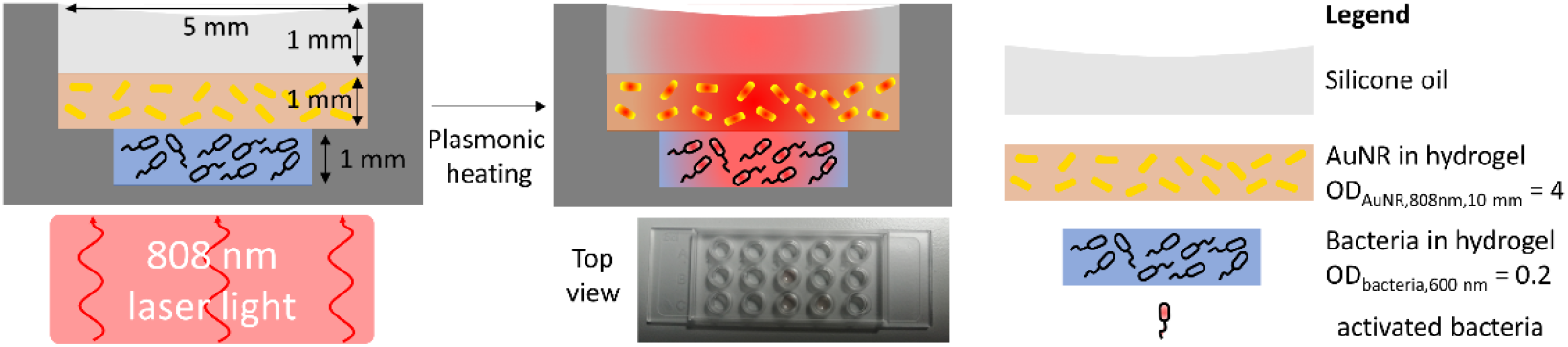
Schematic depiction of the bilayer hydrogel construct in a polymer well plate. The top layer contained the plasmonically absorbing gold nanorods that convert irradiation at 808 nm into heat. This increased the temperature of the bottom layer, which contained bacteria genetically modified to produce mCherry at temperatures above 39°C.

### 2.5 Laser setup

The laser setup for the photothermal excitation of the bilayer hydrogel is schematically depicted in Fig. S1. A continuous wave (CW) infrared laser (CNI, China) emitted light at 808 nm with adjustable power from 1–6 W. The laser light was coupled into a SMA 905 multimode fiber (400 μm core diameter, CNI, China) connected to a collimator (FOC-01, CNI, China) that parallelized the outcoming light to a 5 mm diameter spot. We used neutral density (ND) filters (NE10A-B, Thorlabs, Germany) between collimator and photodiode to reduce outcoming laser power to the range of 100 mW when using laser power of 1 W. A mechanical shutter with an integrated Si photodiode (FDS10×10, Thorlabs, Germany, behind an ND filter from Thorlabs) was used to monitor the laser power.

A dielectric right angle prism mirror (MRA20-E03, Thorlabs, Germany) guided the light to the sample that was at a distance of 10 cm from the collimator inside the patterned microscopy slide (see above). The lateral position of the beam spot on the sample was adjusted with the aid of a monochromic camera (DMK 23U445, The Imaging Source, Germany). An infrared camera (VarioCam@HD 980 S, Infratech, Germany) was mounted 10 cm above the sample to measure the temperature distribution.

### 2.6 Photothermal Conversion Efficiency (PCE)

The photothermal conversion efficiency (PCE) of GNC hydrogels was quantified using published methods.[47, 48] Hydrogels with and without nanorods were exposed to light, the temperature changes were measured and interpreted by balancing the photothermal heat produced by the gold nanorods (*Q*_in_), transient heating of quartz cuvette-hydrogel system (*Q*_0_), and heat dissipation to the external environment (*Q*_out_):[47]

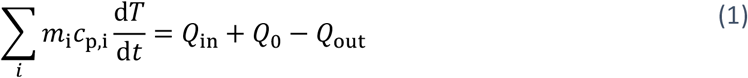

with *m*_i_ and *c*_p,i_ the mass and specific heat capacity of component *i*, and *T* the temperature of all components (assumed to be uniform) at time *t*. Both *m*_i_ and *c*_p,i_ can be determined experimentally. The photothermal conversion efficiency η at steady-state is

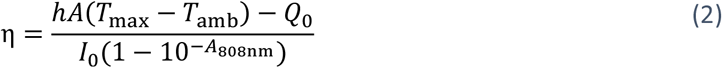

with *h* the heat transfer coefficient, *A* the area of the interface between GNC hydrogel and the external environment, *T*_amb_ the measured ambient temperature, *T*_max_ the measured maximal temperature reached under illumination at steady-state, *I*_0_ the incident laser power, and *A*_808 nm_ the absorbance of the GNC hydrogel at the laser wavelength (808 nm).

To calculate *hA*, we analyzed transient temperature changes. The parameter *B* ≡ *hA*/*m*_i_*c*_p,I_ quantifies the heat dissipation from the GNC hydrogel to the external environment; we assume that it is time-invariant. It can be calculated using an exponential fit of the cooling curve as the laser is turned off:[48]

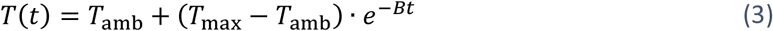

The Supporting Information provides further details on the fit and the PCE calculation.

### 2.7 Temperature measurements for PCE quantification and stability testing

A thermocouple (K-Type, 1.4541 stainless steel) was inserted in GNC hydrogel (OD_AuNR,2mm_ = 0.67, 100 μL) in a glass cuvette (Hellma, Germany) with a beam path length of 2 mm and a glass thickness of 0.5 mm. An infrared diode laser (808 nm, 89 mW) with beam diameter of 10 mm (collimator FOC-01, CNI, China) was used to illuminate the nanorod-containing gel and a reference gel without rods. The temperature change was recorded at 5 s intervals using a 4-channel data logger (TC direct, Germany) via the *SE521* software. The laser was switched off after 10 min and the cooling curve was recorded during the subsequent 10 min. The ambient temperature was measured with an additional TC.

The steady-state temperature profiles in hydrogel bilayers were quantified the wells of the patterned polymer slide. The photothermal stability of the GNC during prolonged and cyclical excitation was evaluated, too. The bilayers were prepared as explained in Fig. 1, but without bacteria in the bottom layer. Top layers with an OD_AuNR,808nm_ = 4 were illuminated with laser power densities of 0.4 W/cm^2^, 0.5 W/cm^2^, 0.6 W/cm^2^, 0.7 W/cm^2^ and 0.8 W/cm^2^; top layers with OD_AuNR,808nm_ = 0, 1, 2, 3, 4, 5 were illuminated with laser power density of 0.8 W/cm^2^ to provide systematic data. Local temperatures at the (oil-air) interface were obtained through infrared thermography using an infrared camera (Infratec, Germany).

### 2.8 Photothermally triggered protein expression

Bilayered ELMs were prepared as explained in Section 2.4 and Fig. 1 with a top layer AuNR concentration equivalent to OD_AuNR,808 nm_ = 4. They were irradiated at laser power densities of 0.5– 0.7 W/cm^2^ to reach different maximal steady-state temperatures during 0 h, 1 h, 2 h, 3 h, and 4 h.

The constructs were then observed under a Keyence BZ-X800 fluorescence microscope with an 4x objective. The magnification was chosen to depict the entire microslide well in one image. We used the filter cubes BZ-X Filter OP-87764 with an excitation of 545 nm (25 nm bandpass) and an emission of 605 nm (70 nm bandpass) at an integration time of 1.5 s of the digital camera. The fluorescence intensities of the acquired images were evaluated via *ImageJ*. A rectangular selection with length 4 mm and height 0.17 mm was chosen to plot profiles for all the fluorescence images. The rectangular selection was rotated by 34.6 °C to evaluate the diagonal of the fluorescence images, thus covering the entire diameter of the well (see Fig. S2). Only the lateral positions between 0.2–3.8 mm were considered to avoid optical scattering effects of the well border.

The lateral temperature distribution during the experiment was quantified by IR thermography and the images were analyzed using the software *IRBIS 3.1*. Steady-state temperature distributions obtained at *t* = 60–595 s were converted to TIFF raw data. A rectangular area with a length of 4 mm, a height of 0.17 mm, and a rotation angle of 34.6° that coincided with that used in fluorescence analysis was analyzed using *ImageJ* to obtain averaged linear profiles.

Each experiment was repeated 3 times and the average local fluorescence intensities and average lateral steady-state temperatures with their standard deviation were plotted.

## 3. Results and Discussion

### 3.1 Preparation and characterization of GNC hydrogels

The Photothermal Conversion Efficiency (PCE) and the uniformity of the GNC hydrogels depends on the agglomeration state of the nanorods. We used commercial, CTAB-stabilized gold nanorods (41 nm long, 10 nm diameter) in aqueous suspension (AuNR@CTAB). They were adjusted to the desired concentrations by centrifugation, mixed with PLUDA_MQ_ at 4° C below the lower critical solution temperature (LCST) of 15 °C,[45] incubated to initiate gelation at room temperature, and cross-linked with UV light. The overall process entails several risks of agglomeration: the concentration of the ligand, CTAB, is lowered by a factor 40 to 0.125 mM during centrifugation due to surfactant removal, possible causing desorption of the weakly bound molecule from the rod. Tebbe *et al*. and Reiser *et al*. reported that gold nanorods are stable at CTAB concentrations of 0.1 mM and 0.08 mM for a short time.[33, 49] On the other hand, AuNR concentration is increased to approximately OD_AuNR, 808 nm_ = 40 during centrifugation. The combination of high AuNR concentration and low concentrations of CTAB increases the agglomeration risks.

We prevented agglomeration by minimizing the time between concentration increase and gelation of the hydrogel, which prevents diffusion,[45] and used optical spectroscopy to assess the agglomeration state in the gels. Pure hydrogels (without AuNR) had an OD of below 0.13 between 400 and 900 nm, with an OD = 0.09 at 808 nm (see Fig. S3), the wavelength that we used for the photothermal heating in this work. The UV-VIS spectrum of an AuNR-containing GNC in Fig. 2(a) shows the localized surface plasmon resonance (LSPR) of an aqueous AuNR dispersion with the transversal (T-LSPR) resonance at 510 nm and the longitudinal resonance (L-LSPR) at about 808 nm. The L-LSPR peak shifted from 805 nm to 803 nm in the gel, as expected due to the 30 wt-% of PLUDA polymer that increases the refractive index of the rods’ environment. Note that the L-SPR is more sensitive to the environment than the T-SPR due to the larger longitudinal polarizability,[16, 50] as visible in the plot. The full width half at maximum (FWHM) of the L-LSPR changed from 111.3 ± 0.4 nm for AuNR@CTAB to 118.6 nm ± 0.4 nm in the gel, indicating no or very limited agglomeration.[51, 52] This is consistent with cryo-TEMs (Fig. 2(b)) that show well-distributed rods.

**Figure 2:**
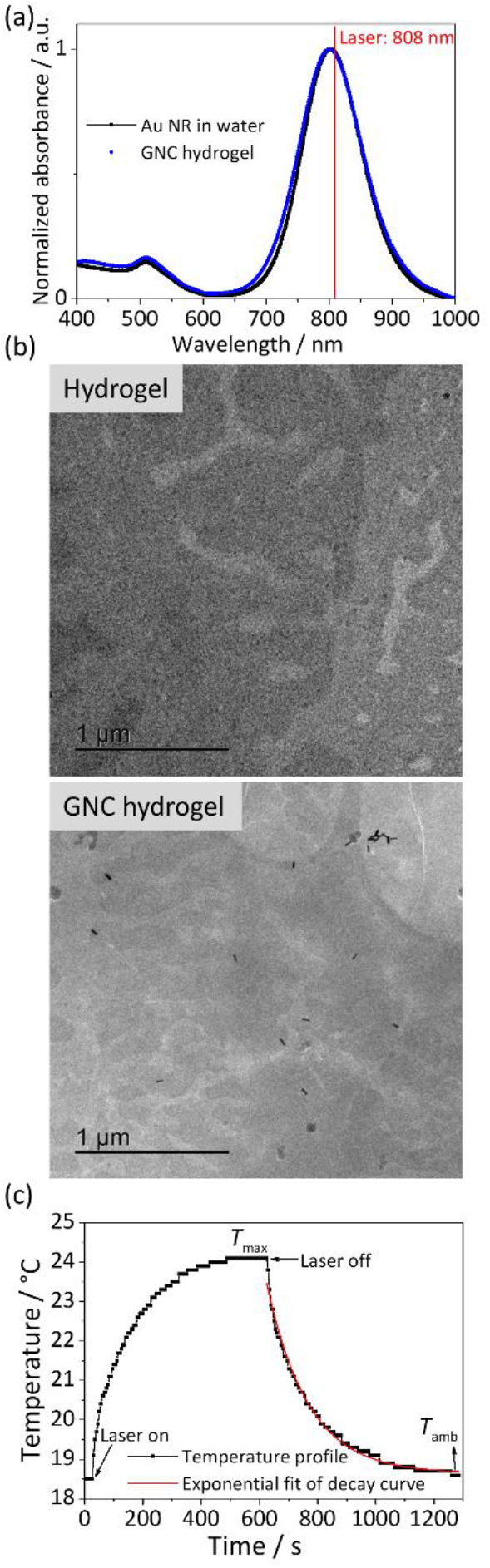
Agglomeration state of the AuNR in the GNC gel. (a) Normalized UV-VIS absorbance spectra of an aqueous dispersion of AuNR@CTAB and the GNC hydrogel. (b) Cryo-TEM of the pure hydrogel (top) and the GNC hydrogel (bottom). (c) Typical heating and cooling profile of a laser illuminated GNC hydrogel. The red line represents the exponential decay fit of Eq. (3).

Nanorods with a L-LSPR at 808 nm were chosen to match the transmissive window of tissues and ensure the availability of excitation lasers. In the following, we analyze whether the photothermal conversion efficiency of the GNC hydrogels is comparable to those reported for GNC dispersions using an established measurement technique reported by Roper *et al*.[47] A high PCE is critical to avoid exceeding the irradiation limit of 0.3 W/cm^2^ for 808 nm lasers on human tissue.[53] Typical mammalian tissues attenuate 808 nm light to 10 % of the incoming intensity in 1 cm.[54]

Fig. 2(c) shows a typical heating curve of an illuminated GNC hydrogel with a maximal temperature increase of 5.7 ± 0.11 °C at a laser power of 0.09 W and an OD_2mm,808nm_= 0.67. The steady state was reached after approximately 600 s. Cooling after laser deactivation was exponential and reached *T*_amb_ = 18.6 °C after 600 s. It was used for the quantification of the PCE[47, 48, 55] by fitting Eq. (3) that indicated a heat dissipation rate *B* = 0.00693 ± 8·10^−5^ s^-1^ and thus, a PCE of η = 47 ± 0.01 % from Eq. (2). This is the first reported value of a PCE of gold nanorods in a gel to the best of our knowledge. The value, an intrinsic property that depends on the shape and size of AuNRs,[55, 56] is comparable to that from previous reports on AuNR in water of 50 % and 55 %.[57, 58]

### 3.2 Photothermal activity and stability of the bilayer hydrogel

We prepared bilayers in microwells with pure hydrogel as bottom and GNC hydrogel as top layer. The bottom layer will be loaded with encapsulated bacteria in later experiments; we assume that they do not significantly affect the photothermal properties of the gel because their volume fraction is low (< 8 % v/v according to literature[45]) and their heat capacity close to that of the gel. The bilayers were formed by adding the liquid gel precursors at 4 °C. They gelled at room temperature and were stabilized by photochemical UV cross linking.

Illumination of the bilayer hydrogel with 808 nm light resulted in heating. The gold nanorods in the top layer absorbed the light and converted it into heat. Fig. S4 shows the relevant heat flows: heat is thermoplasmonically generated at a rate *Q*_gen_ in the top layer. It diffuses to the surroundings, heating the lower layer by heat conduction *Q*_cond,1_ and the silicon oil by conduction *Q*_cond_ and convection *Q*_conv_. Some heat is lost by diffusion through the polymer microwells, *Q*_cond,2_. Convection *Q*_air,conv_ to the surrounding air finally removes heat from the system. In steady state, heat is constantly dissipated to the environment, and a constant temperature distribution in the bilayer hydrogel emerges. We measured the temperature at the silicone-air surface using an infrared camera. Fig. S5 shows representative IR image of the construct.

We quantified the photothermal excitation of the bilayer construct by analyzing the temperature at a defined point of the silicon oil (marked in Fig. S5) as a function of time. When irradiated with 808 nm light, this temperature increased and reached a plateau after approximately 6 min (Fig. 3(a) and (c)). The thermal stability of the construct was tested by repeated illumination (laser power density 0.8 W/cm^2^ and OD_AuNR, 808nm_ = 4) and cooling (by turning the laser off) (Fig. 3(d)) in three cycles. Heating and cooling curves followed the expected exponential kinetics. These results confirm stable and efficient photothermal heating of the hydrogel bilayer.

**Figure 3:**
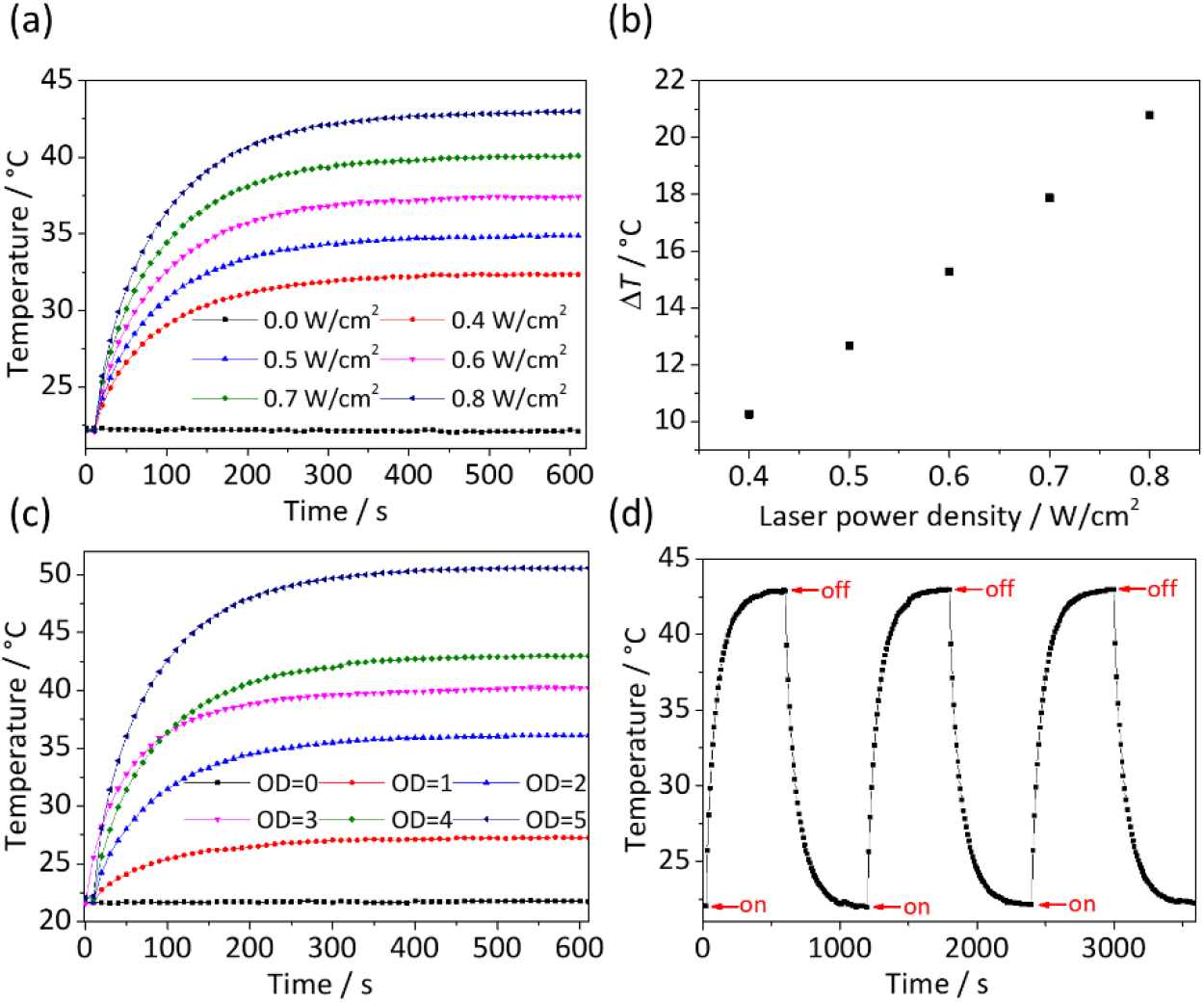
Photothermal heating of bilayer structures. (a) Heating of illuminated hydrogel bilayers at a GNC OD_AuNR,808nm_ = 4 and different laser power densities. (b) Plateau temperature differences of the GNC as a function of laser power density. The differences between the initial temperatures of the hydrogels and the maximal steady state temperatures are shown. (c) Heating of illuminated hydrogel bilayers at 0.8 W/cm^2^ for different GNC OD. (d) Stability test by intermittent illumination of a GNC with OD_AuNR,808nm_ = 4 at a laser power density of 0.8 W/cm^2^.

In steady-state, *Q*_gen_ equals the dissipating heat output *Q*_cond,i_ and *Q*_conv._ At a laser power density of 0.8 W/cm^2^ and AuNR concentration OD_AuNR,808nm_ of 4, a steady-state difference to the environmental temperature of Δ*T* ≈ 22 °C had stabilized after 6 min. We varied the incident laser power density from 0.4 W/cm^2^ to 0.8 W/cm^2^ at an AuNR concentration of OD_AuNR, 808nm_ of 4 and found Δ*T* between 10 °C and 21 °C in accordance with Eq. (2) (Fig. 3(b)). Increasing AuNR concentrations OD_AuNR, 808nm_ from 1 to 5 at a laser power of 0.8 W/cm^2^ led to Δ*T* between 5 °C and nearly 29 °C (Fig. 3(c)).

### 3.3 Temperatures inside the bottom layer of the bilayer hydrogel

Infrared thermometry provided the temperature at the silicone oil surface. We were interested in the temperatures inside the bulk gel and embedded a thermocouple (TC) within the bottom hydrogel layer. Fig. 4(b) shows a representative surface temperature profile at 0.6 W/cm^2^ from the IR camera and a plot of the surface temperature *T*_IR_ at position P1 (Fig. 4(c)). The same plot shows the temperature concurrently measured inside the gel with a TC (*T*_TC_) directly below P1 at a depth of 3 mm.

**Figure 4:**
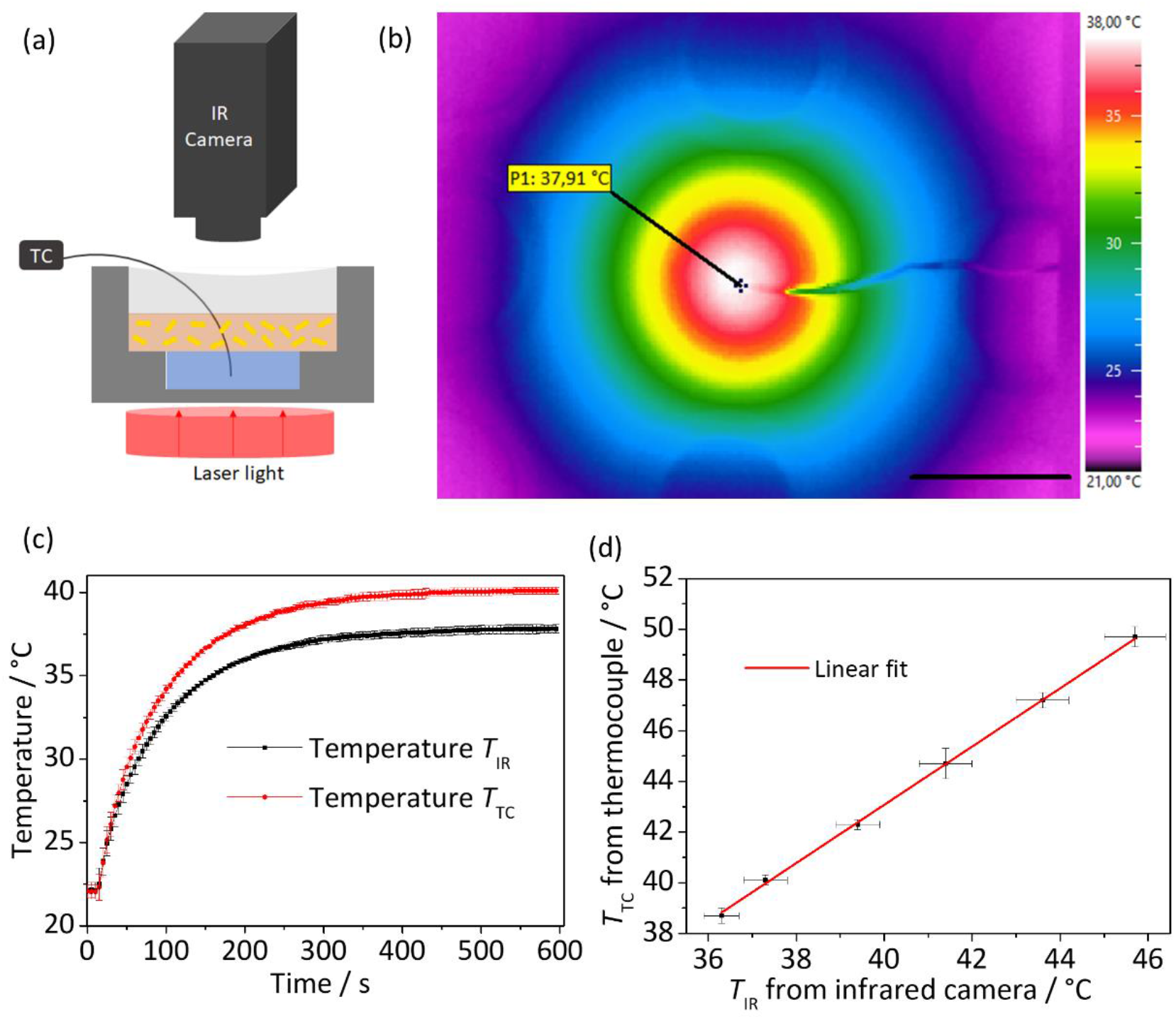
Comparison of surface and bulk temperatures. (a) Cross section of one well of the microslide depicting the TC and IR camera position. (b) Steady-state temperature profile from IR thermometry at a laser power density of 0.6 W/cm^2^ and OD = 4. P1 denotes the position used for comparing IR measurements with the reading from an embedded thermocouple (TC). The scale bar is 5 mm. (c) Temperatures T_IR_ measured at P1 by IR and T_TC_ measures below P1 by TC during transient heating at 0.6 W/cm^2^ (average of three measurements, provided with standard deviation) (d) Relation of T_TC_ and T_IR_ for steady-state temperatures. The steady state temperatures at t = 595 s were averaged, and their standard deviation calculated at laser power densities of 0.57 W/cm^2^, 0.62 W/cm^2^, 0.70 W/cm^2^, 0.77 W/cm^2^, 0.85 W/cm^2^ and 0.93 W/cm^2^.

The temperatures inside the gel were always above the surface temperature. We measured steady state temperatures for laser powers ranging from 0.6 W/cm^2^ to 1.0 W/cm^2^ at an OD_AuNR,808nm_ of 4 and found a linear correlation between *T*_TC_ and *T*_IR_. This enables a simple conversion of the IR observation into bulk temperature for point P1:

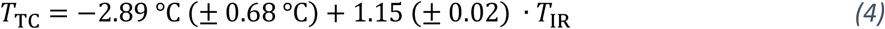

Note that while the precision of the measurement is good, its accuracy is limited by the positioning of the TC and the reproducibility of the hydrogel layer geometry. We estimate errors for 2 mm of spatial uncertainty (see Fig. S7) for the positioning of the TC. This corresponds to uncertainties of the measured temperature on the order of 0.4–0.7 °C for laser power densities of 0.57–1.0 W/cm^2^. Note also that there exist both lateral and normal temperature gradients in the gel such that bacteria experience different temperatures depending on their exact position. We tuned laser power in all subsequent experiments such that the bottom hydrogel was in a range of 39–50 °C and report the surface temperatures of the IR camera that can be converted into bulk temperatures using Eq. 4.

### 3.4 Thermoplasmonic NIR stimulation of mCherry producing thermoresponsive bacteria

We then tested the ability of the gold nanorod containing hydrogels to activate ClearColi bacteria that were genetically engineered to thermoresponsively express a red fluorescent protein (mCherry). The bacteria were engineered with a TlpA regulator[46] that had a sharp switch-like transition from keeping gene expression OFF at 37 °C to switching it ON above 39 °C (See Fig. S8). The bacteria were encapsulated in the bottom layer and covered by a GNC hydrogel layer (Fig. 1). The temperature of the surface layer (silicon oil) was determined from IR thermography. Power densities of 0.5–0.7 W/cm^2^ of the 808 nm laser were set to achieve surface temperatures between 35 °C and 41.5 °C. The bottom layer temperature can be estimated from this temperature using Eq. 4; it is always above the surface temperature and was sufficient to activate the bacteria.

Activation of the thermoresponsive bacterial switch was indicated by the expression of mCherry, whose fluorescent signal across the entire bacterial hydrogel was detected using fluorescence microscopy initially and after each hour four maximal 4 hours of photothermal stimulation. Such microscopy imaging allowed spatial quantification of fluorescence intensities with a spatial resolution down to 2 μm in the XY plane (Fig. 5(a)–(b)). Fig. S9(a)–(b) shows the increase of fluorescence intensity with photothermal stimulation time. A uniform expression of mCherry was observed after photothermal heating at 0.7 W/cm^2^ laser power density, equivalent to a temperature range of 38– 41.5 °C on the silicon oil surface depending on the lateral position and a higher temperature in the bulk gel (Fig. 5(d)). We conclude that this level of heating created an ideal steady-state temperature range to achieve the highest rate of gene expression while keeping the bacteria viable. Heating with 0.5– 0.6 W/cm^2^ led to a surface temperature profile in a range of 35–37.5 °C, leading to maximum mCherry expression in the center that dropped to low levels at the periphery as visible Fig. 5(a) and (c). This indicates the possibility to achieve locally confined activation of ELM functions using this system. Thus, by tuning the laser power at same GNC concentrations, it is possible to influence the spatial profile in which the ELMs can be activated.

**Figure 5:**
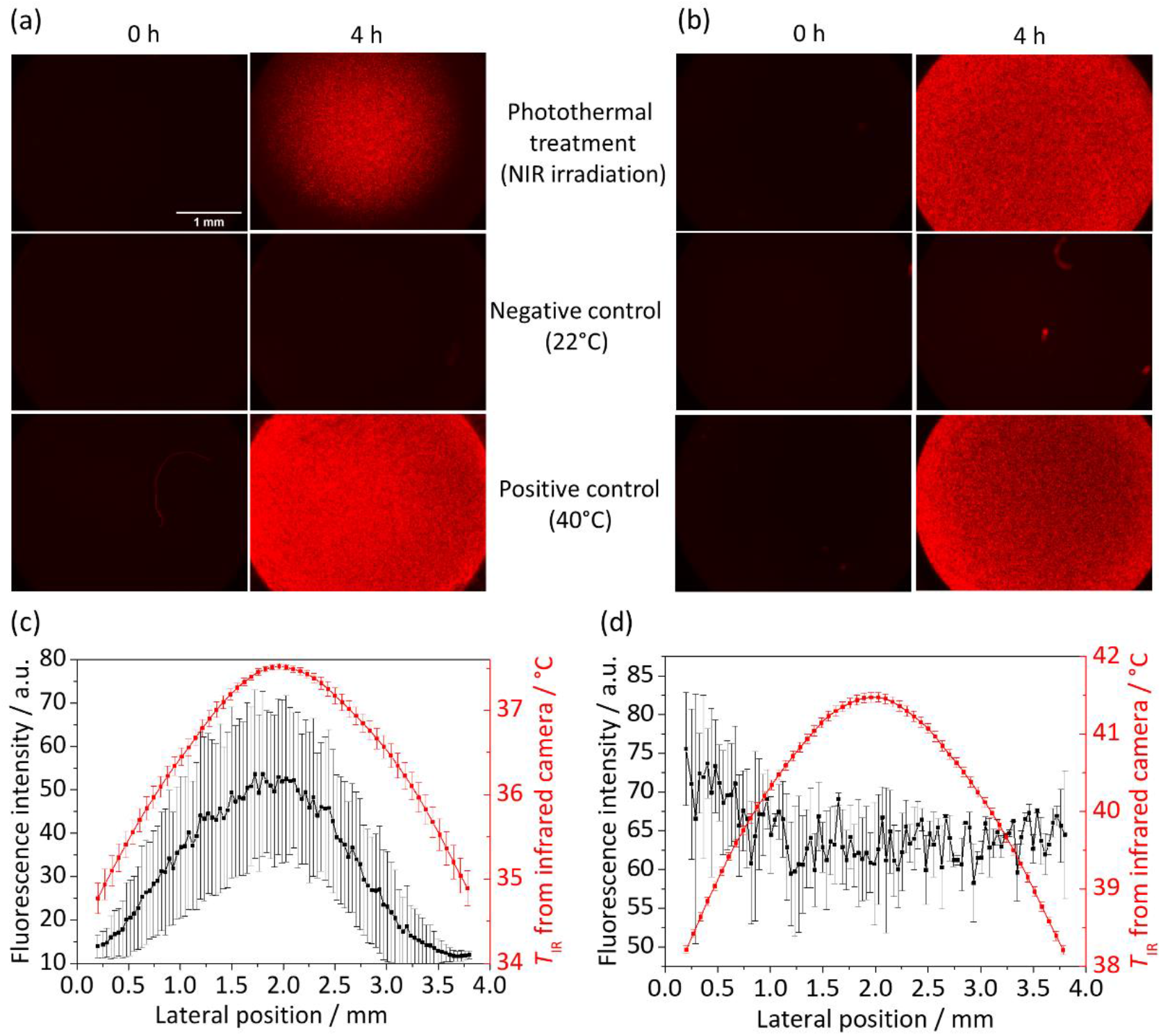
Expression of fluorescent mCherry in thermoplasmonically stimulated ELMs. Fluorescence micrographs of ELM before and after 4 h stimulation at 0.5 W/cm^2^(a) and (b) 0.6 W/cm^2^. The temperatures reached in (a) were sufficient to activate bacteria in the center, but not at the border. The temperatures reached in (b) were sufficient for uniform activation, leading to uniform fluorescence. (c) Mean fluorescence and surface temperature profiles for laser power densities of 0.5–0.6 W/cm. (d) Profiles for a laser power density of 0.7 W/cm^2^.

The stability of the steady-state temperature profiles was excellent, with deviations below 1 K over several hours at each of the individual positions. Variations between different samples were due to uncertainties in the exact AuNR concentration of the GNC and the geometry of the casted gels. Pipetting identical Pluronic precursor solutions at low volumes is difficult since they quickly form physical gels above 10 °C and form menisci at the walls. This explains deviations that we found e.g. between the three different samples shown in the error bars of Fig. 5(c))–(f). Such uncertainties can be reduced by improved processing when larger amounts of ELM are prepared.

## 4. Conclusion

We designed an ELM with a bilayer hydrogel structure containing AuNRs and thermoresponsive bacteria and demonstrated the use of NIR light for the activation of the bacteria through photothermal conversion. This provides new options for the on-demand activation of ELM activity for therapeutic applications.

We demonstrated that the colloidal stability of AuNR@CTAB in hydrogel is sufficient to prepare NIR absorber gels with high efficiency. Agglomeration was not observed, and reproducible photothermal heating was achieved with the GNC hydrogel. The PCE of the GNC hydrogel was around 47 %, comparable with literature reports on pure nanorods, sufficient for the application, and stable over multiple cycles. Illumination with a collimated laser spot of 5 mm diameter at 808 nm and a power density of 0.6–0.7 W/cm^2^ heated a cylindrical bilayered hydrogel in air from an initial 22 °C to a steady-state temperature of 41 °C in 300 s. Infrared thermometry indicated a lateral surface temperature gradient from center to border with a maximum temperature difference of 4.7 °C.

The irradiation limit on human tissue (maximum permissible exposure, MPE) equals 0.3 W/cm^2^ for 808 nm lasers.[53] Light at 808 nm penetrating mammalian tissues was attenuated to 10–20 % at 1 cm.[54] Assuming a linear relation between generated temperature difference and laser power density,[47] and the data reported above, we would expect a temperature increase on the order of 1 K for the temperature increase of a Living Material embedded in tissue at 1 cm depth at a laser power density of 0.3 W/cm^2^. The true value will be considerably larger because the convective cooling in our experiments is more efficient than the thermal diffusion that will remove heat from the implanted ELM. We are currently working on experimental setups to better emulate this situation. In addition, it will be possible to increase the temperature at lower laser power densities of 0.3 W/cm^2^ by increasing the concentration of gold nanorods; one order of magnitude in concentration is probably attainable with suitable ligands that prevent agglomeration. An additional route to larger temperatures is a switch of laser wavelength to 1275 nm with an MPE of 1 W/cm^2^ and higher penetration depths in tissue,[59] at the cost of more expensive laser sources and optics.

We quantified bacterial activity in a GNC-sensitized ELM as a function of laser power, time, and position via *ex situ* fluorescence microscopy. The fluorescence intensity correlated with the temperature distribution. Protein production at 0.5–0.6 W/m^2^ was maximal at the center of the illuminated area and declined at its borders, where temperature dropped. Increasing laser power first led to an increase of fluorescence intensity at the borders. A uniform fluorescence intensity was reached across the illuminated area at 0.7 W/m^2^. Protein expression continued for at least 4 h, with a linear increase of fluorescence that indicated sustained expression activity of the bacteria.

In summary, we demonstrated the NIR stimulation of an ELM and its efficacy using a model bacterium. This concept can be expanded to include additional stimuli such as magnetic fields or ultrasound in order to create multi-stimulus ELMs.[60, 61]

## Acknowledgement

Funding from the Leibniz Science Campus “Living Therapeutic Materials” is gratefully acknowledged. The authors thank Marcus Koch for his support in cryo-TEM imaging of the GNC and Peter Rogin and Robert Strahl for fruitful discussions about the laser setup. The [pTlpA39-Wasabi](https://www.addgene.org/86116/) vector was a gift from Prof. Mikhail Shapiro, Addgene #86116.

