## Supporting information for "Plasmonic Stimulation of Gold Nanorods for the Photothermal Control of Engineered Living Materials"

### Laser setup

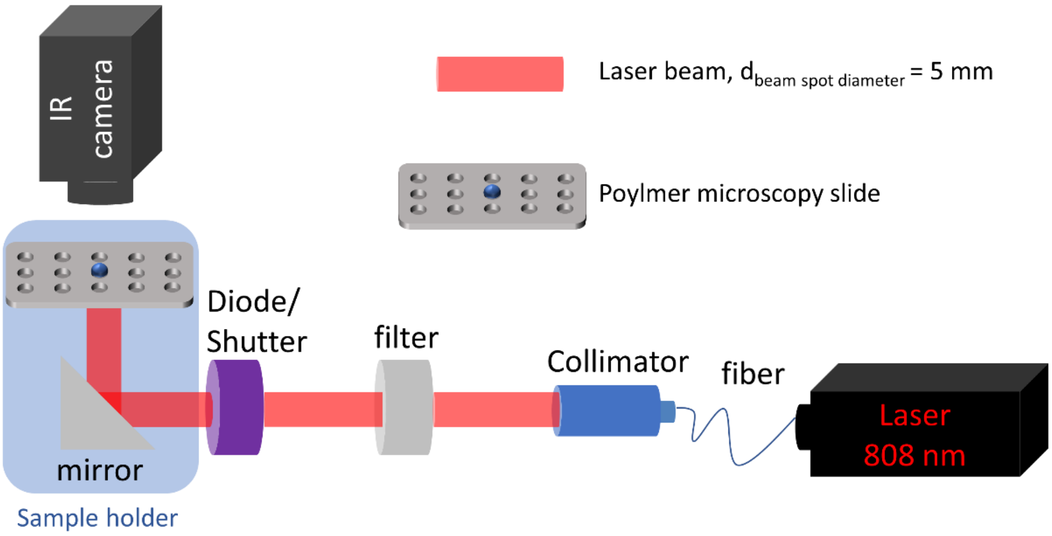


*Figure S 1: Laser setup used for laser excitation at 808 nm with concurrent observation of surface temperature with an IR camera.*

### Fluorescence intensity analysis in micrographs

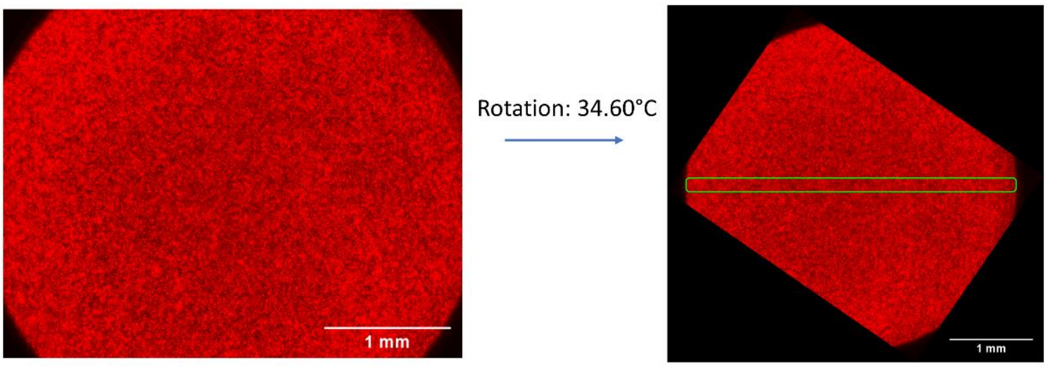


*Figure S 2: Quantification of fluorescence intensity via image analysis. The fluorescence micrograph was analyzed using Image J by rotating the image by 33.6° and inscribing a rectangle with 0.17 mm height and 4 mm length. The pixel values were then averaged along the height and the mean value used as a line profile.*

#### UV–VIS absorbance spectra from hydrogel

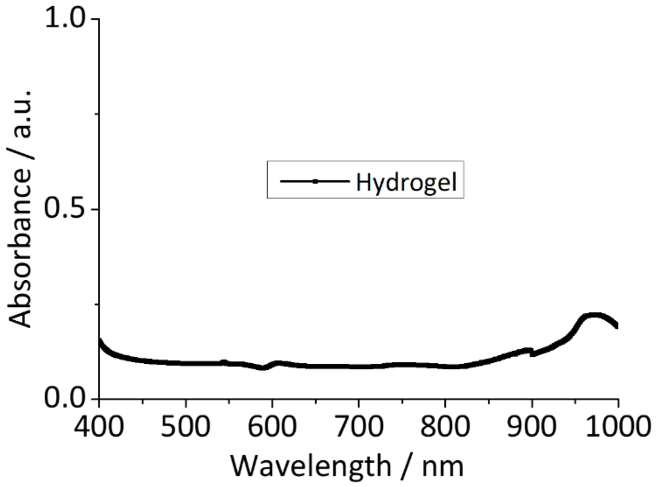


*Figure S 3: UV–VIS spectra from hydrogel without embedded Au NR.*

#### Heat transfer in hydrogel bilayer in a microwell

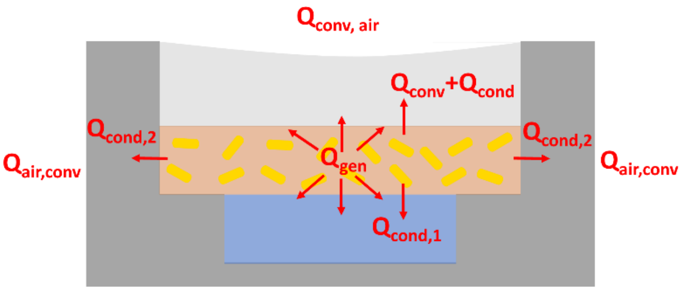


*Figure S 4: Schematic structure of the hydrogel bilayer in a microwell. Schematic structure of the hydrogel bilayer in a microwell. Thermal transport dissipated the heat generated photothermally at a rate $Q_{gen}$ through thermal diffusion and convection to the environment. The result was a steady-state temperature distribution at the surface that we quantified.*

#### Position P1 for evaluation of photothermal activity and stability

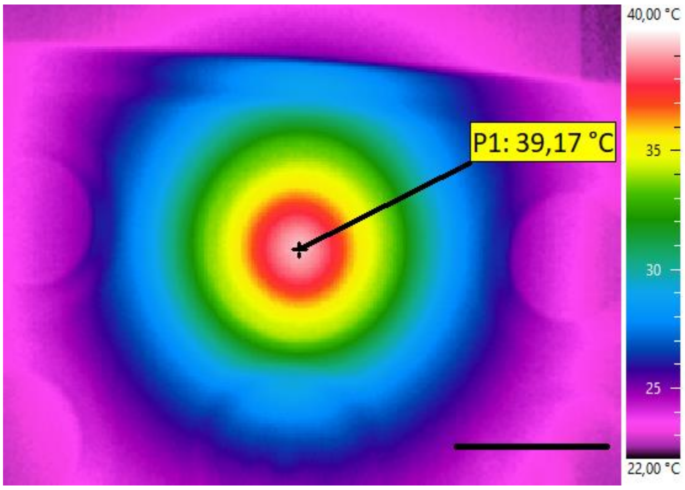


*Figure S 5: Infrared thermography image with the marked position P1 that we used for the photothermal stability measurements in section 3.2. Representative steady state temperature shown here. Scale bar indicates 5 mm.*

**Setup for determing Photothermal Conversion Efficiency (PCE)**

The PCE of the gold nanorod composite (GNC) hydrogel was determined in a glass cuvette. A thermocouple was embedded in the GNC hydyrogel to follow the temperature changes induced by thermoplasmonic heating upon 808 nm laser irradiation.

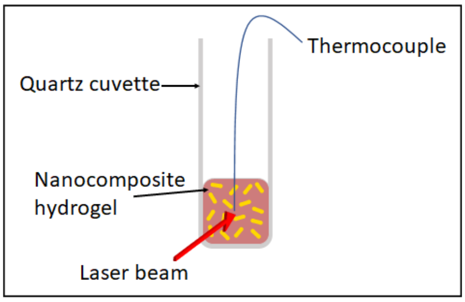


*Figure S 6: Setup for the determination of the PCE of a gold nanorod composite (GNC) hydrogel in a glass cuvette. The thermocouple was used to quantify the temperature change upon illumination.*

**Calculation of the Photothermal Conversion Efficiency**

The photothermal conversion efficiency was calculated following literature.[1-4] The total energy balance of the system is described as:

$$\sum_i m_\mathrm{i} c_\mathrm{p,i} \frac{\mathrm{d}T}{\mathrm{d}t} = Q_\mathrm{in} + Q_0 - Q_\mathrm{out} \tag{1}$$

where $m_i$ and $c_{p,i}$ are the mass and heat capacity of GNC hydrogel and glass cuvette in the system, $T$ is the system temperature, and $t$ is time.
$Q_{in}$ is the photothermal energy emitted by the illuminated AuNRs under laser light illumination:

$$Q_\mathrm{in} = I_0 (1 - 10^{-A_{808\mathrm{nm}}}) \eta \tag{2}$$

where $I_0$ is the incident laser power (89 mW), $A_{808nm}$ is the absorbance of GNC hydrogel in cuvette at 808 nm which was measured to be $OD_{AuNR,2mm,808nm}$ = 0.67, and η is the photothermal conversion efficiency. $Q_0$ is the energy input contributed to adsorption by the pure hydrogel and the cuvette and was determined to be 0.35 mW.
The heat dissipated to the external environment is given by $Q_{out}$:

$$Q_{out} = hS(T(t) - T_\mathrm{amb}) \tag{3}$$

where $h$ is the heat transfer coefficient, $S$ is the surface are of the quartz cuvette, $T(t)$ is the temperature of the GNC hydrogel at time $t$, and $T_{amb}$ is the ambient temperature of the surroundings.
By defining $\Delta T \equiv T(t) - T_{amb}$, equation 1 can be expressed as:

$$\frac{d\Delta T}{dt} = \frac{I_0 (1 - 10^{-A_{808\mathrm{nm}}}) \eta}{\sum_i m_\mathrm{i} c_\mathrm{p,i}} - \frac{hS}{\sum_i m_\mathrm{i} c_\mathrm{p,i}} \Delta T + Q_0 \tag{4}$$

A time constant of the overall system is defined as $B \equiv hS / \sum m_i c_{p,i}$. It is determined experimentally by measuring the decreasing temperature profile after the laser is turned off, setting $Q_{in} = 0$ in equation 4, and setting $T(t = 0\ \mathrm{s}) = T_{max}$, resulting in the cooling temperature profile:

$$T(t) = T_\mathrm{amb} + (T_\mathrm{max} - T_\mathrm{amb}) e^{-Bt} \tag{5}$$

where $T_{max}$ is the maximum temperature at which the laser is turned off.
The cooling curve was fitted with Eq. 5 to determine the constant $B$. We use it to calculate $hS$, making it possible to determine the PCE η:

$$\eta = \frac{hA(T_{\max} - T_{\mathrm{amb}}) - Q_0}{I_0(1 - 10^{-A_{808\mathrm{nm}}})} \tag{6}$$

**IR image for determing the lateral uncertainty of section 3.3**

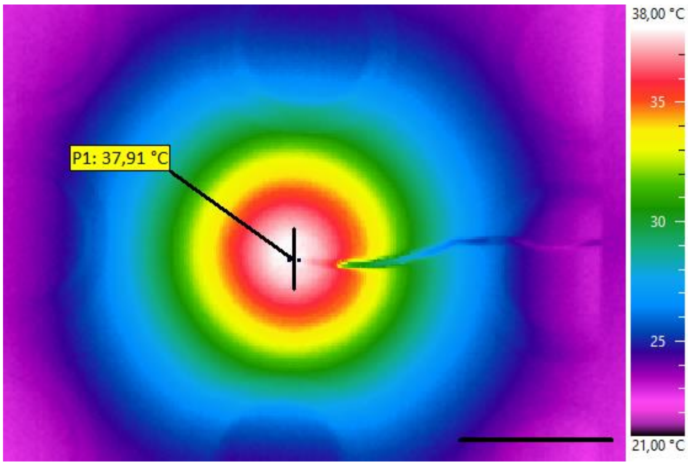


*Figure S 7: Infrared thermography image of the steady state of a illuminated, bilayered hydrogel. The inset at the center (length: 2 mm) indicates the temperature gradient that we used to calculate the lateral uncertainty in temperature. Scale bar indicates 5 mm.*

**Temperature range for the bacterial activity**

Bacterial cultures were incubated overnight in 5 mL of LB-NaCl media (supplemented with 100 µg/mL ampicillin) at 30°C with continuous shaking (250 rpm). The following day, cultures were diluted to 0.1 $OD_{600}$ in 3 mL of antibiotic-supplemented fresh media and regrown at 30°C, 250 rpm. At $OD_{600}$ = 0.3, the cultures were dispensed into Fisherbrand™ 0.2mL PCR Tube Strips with Flat Caps (Thermo Electron LED GmbH, Germany) and placed in the Biometra Thermocycler (Analytik Jena. GmbH, Germany). The thermal assay was set at a temperature gradient from 31◦C to 43◦C with regular increment of 2◦C. The lid temperature was set at 50◦C to prevent the evaporation of the liquid and maintain a homogeneous temperature in the spatially allocated PCR tubes. After a time interval of 18 h, the PCR strips were centrifuged in a tabletop minicentrifuge (Biozym GmbH, Germany) to pellet down the cells and discard the supernatant. The cells were then resuspended in 200 µL of 1X PBS and added to the clear bottom 96-well microtiter plate (Corning® 96 well clear bottom black plate, USA). The samples were then analyzed in the Microplate Reader Infinite 200 Pro (Tecan Deutschland GmbH, Germany) and both the absorbance (600 nm wavelength) and mCherry fluorescence intensity ($Ex_\lambda$ / $Em_\lambda$ = 587 nm/625 nm) were measured. The z-position and gain settings for recording the mCherry fluorescent intensity were set to 19442 µm and 136 respectively. Fluorescence values were normalized with the optical density of the bacterial cells to calculate the Relative Fluorescence Units (RFU) using the formula RFU = Fluorescence/$OD_{600}$.

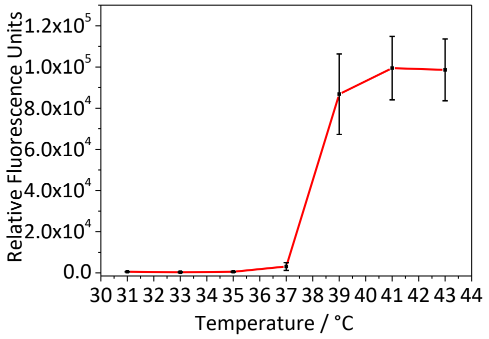


*Figure S 8: Fluoresence intensity of mCherry produced by the Clearcoli bacteria expressed relative fluorescence units as a function of temperature. Symbols represent means and whiskers represent standard deviation from three individual experiments.*

### Thermoplasmonic NIR stimulation of mCherry producing bacteria

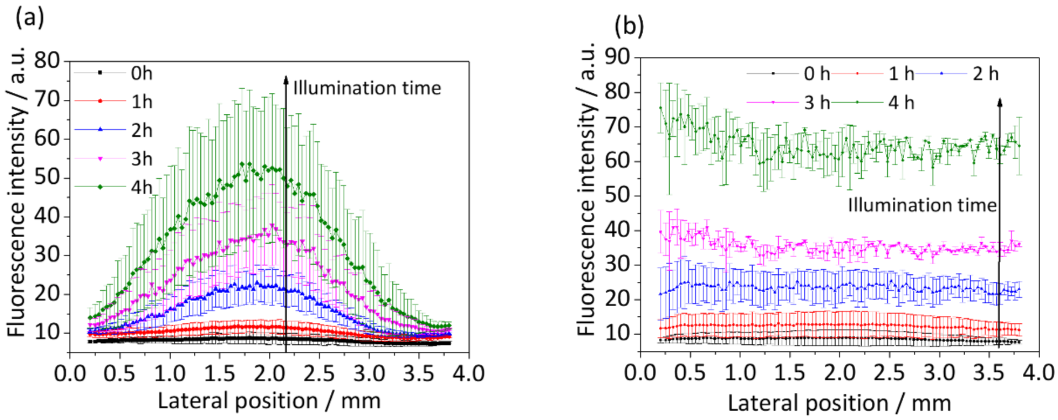


*Figure S 9: Expression of fluorescent mCherry in thermoplasmonically stimulated ELMs at illumination times 0 h, 1 h, 2 h, 3 h and 4 h. (a) Mean fluorescence and surface temperature profiles for laser power densities of 0.5–0.6 W/cm. (b) Profiles for a laser power density of 0.7 W/cm².*

**Positive and negative controls for photothermally simulated ELM**

The expression of the mCherry protein upon photothermal stimulation was compared to positive and negative controls. As a positive control, the bilayered hydrogel was incubated at 40 °C in an oven for 0 h, 1 h, 2 h, 3 h, and 4 h. Uniform bacterial activity was observed. As a negative control, the sample was kept at 22 °C. No bacterial activity was observed.

As an additional negative control, non-engineered bacteria ("wild type") were encapsulated in a gel, exposed to NIR Irradiation, and incubated at 40°C. No mCherry production was detected.

Positive control: Negative control:

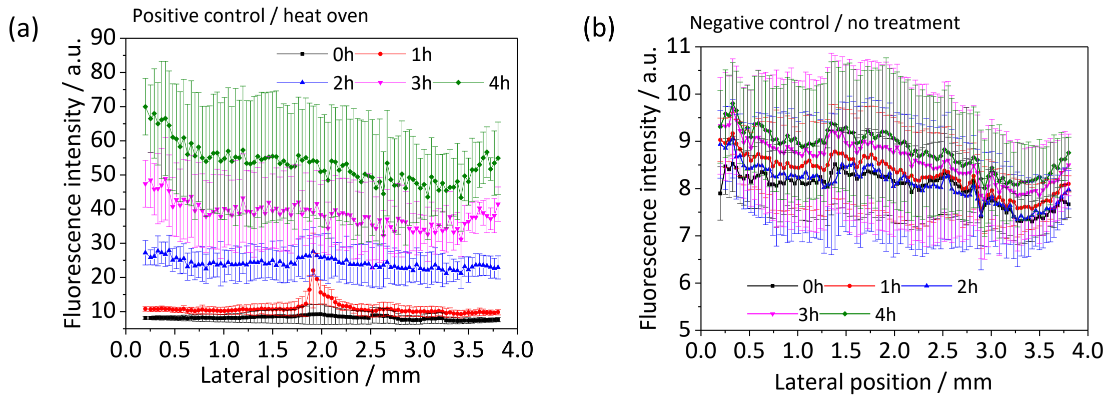


Wild type:

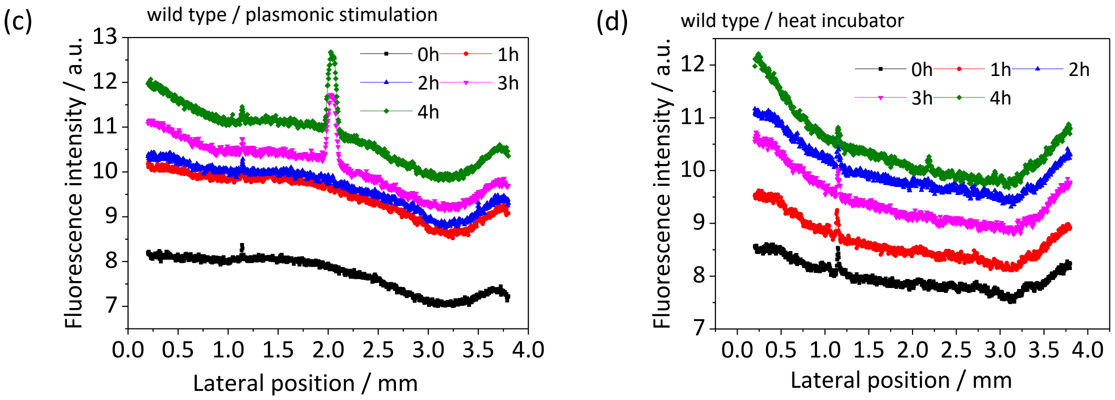


*Figure S 10: (a) The engineered bilayer hydrogel ELM was incubated in heat oven at 40°C as positive control. An increase of mCherry production by continuous time was observed. (b) Negative control of mCherry production of the engineered bilayer hydrogel. The hydrogel was incubated at room temperature at 22°C without any stimulation. No significant mCherry production was observed. (c)+(d) wild type was encapsulated at the bilayer structure. (c) Plasmonic stimulation. The spatial mCherry production was plotted and their fluorescence intensity is shown with different plasmonic duration. No significant mCherry production was detected. (c) Heat incubation at 40°C. No significant mCherry production was observed.*
